# Synergistic and antagonistic activities of *IRF8* and *FOS* enhancer pairs during an immune cell fate switch

**DOI:** 10.1101/2024.09.17.613195

**Authors:** Antonios Klonizakis, Marc Alcoverro-Bertran, Pere Massó, Joanna Thomas, Luisa de Andrés-Aguayo, Xiao Wei, Vassiliki Varamogianni-Mamatsi, Christoforos Nikolaou, Thomas Graf

**Author notes:** Equal contributions (Marc Alcoverro-Bertran, Pere Massó).

## Abstract

Cell-fate instructive genes tend to be regulated by large clusters of enhancers. Whether and how individual enhancers within such clusters cooperate in regulating gene expression is poorly understood. We have previously developed a computational method, SEGCOND, that identifies hubs consisting of enhancer clusters and their target genes, termed Putative Transcriptional Condensates (PTCs). Using SEGCOND, we identified PTCs in a CEBPA-induced B-cell to macrophage transdifferentiation system. We found them to be enriched for highly expressed, lineage-restricted genes and to associate with BRD4, a component of transcriptional condensates. Here we performed single and combinatorial deletions of enhancers within two active PTCs after transdifferentiation is induced, harboring *IRF8* and *FOS*. Two enhancers within the *IRF8* PTC were found to form a backup mechanism when combined, safeguarding *IRF8* expression and transdifferentiation kinetics. Unexpectedly, two individual enhancers within the *FOS* PTC antagonize each other at Day 1 of transdifferentiation, delaying the conversion of B-cells to macrophages and reducing *FOS* expression, but cooperate to increase *FOS* levels in Day 7 induced cells. Our results reveal differentiation stage-specific, complex interactions, between individual enhancers within a cluster.

## Introduction

How cell-type-specific gene expression patterns are established during development and differentiation is a major research topic in current biology (Nora et al., 2023). Evidence accumulated over the past decades has revealed enhancers as major determinants of gene regulation and cell fate determination (Gasperini et al., 2020). Enhancers are sequence-specific DNA elements that recruit transcription factors and co-factors, and that interact with other enhancers as well as with promoters, increasing gene expression. It is well-accepted that large clusters of enhancer elements, described as “LCRs”, “super” or “stretch” enhancers, regulate many cell-fate-specifying genes and drive high levels of gene expression (Grosveld et al., 1987)(Parker et al., 2013)(Whyte et al., 2013). The role of individual enhancers within such large assemblies is not fully understood (Panigrahi & O’Malley, 2021).

Whether individual enhancers within clusters act independently or synergistically is an area of active research (Uyehara & Apostolou, 2023). Some studies suggested that they are essentially independent and act in an additive manner, ensuring that after the loss of one or more enhancers, genes critical for differentiation remain activated (Bahr et al., 2018) (Osterwalder et al., 2018). According to this model, individual enhancers in the cluster may be completely or partially redundant, showing only mild effects when ablated individually. Thus, enhancer clustering could act as a “safeguard” mechanism by conferring expression robustness to their target genes. Other studies, however, have argued that the coalescence of multiple individual enhancers in an assembly may lead to synergistic activatory effects and, consequently, very high expression levels (Hnisz et al., 2017). Indeed, synergistic interactions among individual enhancer elements have been described (Blayney et al., 2023) (Thomas et al., 2021) (Choi et al., 2021). Enhancer synergy has been proposed to be achieved via the formation of transcriptional condensates (Hnisz et al., 2017). Such condensates are membrane-less organelles that contain multiple transcription factors and co-factors such as MED1 and BRD4 and are described to assemble at super-enhancers (Sabari et al., 2018). However, the role of transcriptional condensates is still somewhat controversial (Stortz et al., 2024).

Several computational frameworks have been developed for the genome-wide identification of enhancer clusters (Murphy et al., 2023)(Klonizakis et al., 2023)(Mota-Gómez et al., 2022)(Parker et al., 2013)(Whyte et al., 2013). One of the most widely used is the ROSE algorithm, which detects enhancer clusters termed super-enhancers (Whyte et al., 2013). ROSE stitches together *a priori* identified enhancers and ranks them based on the occupancy of a master transcription factor or a transcriptional co-activator such as MED1 or BRD4. Based on such single ChIP-seq datasets, enhancers that surpass an occupancy threshold are classified as super-enhancers. To improve the detection of enhancer clusters and their target genes, we have previously developed a computational algorithm, SEGCOND, which integrates multiple distinct datasets, including 3D genome conformation data (Klonizakis et al., 2023).

SEGCOND partitions the genome into segments based on chromatin accessibility, histone mark, and transcription factor occupancy data. It then pinpoints segments that are enriched for H3K27ac decorated enhancer elements and, via the integration of Hi-C data, selects those that contain strongly interacting enhancer-enhancer or enhancer-gene pairs. As a final step, SEGCOND links together enhancer-enriched segments of the same chromosome that associate in 3D space (Klonizakis et al., 2023). The final identified regions were originally coined as ‘Putative Transcriptional Condensates’ (PTCs), based on the hypothesis that enhancer clusters serve as scaffolds for the formation of transcriptional condensates (Sabari et al., 2018)(Hnisz et al., 2017). PTCs can thus consist of one, or more genomic segments and contain enhancers and their target genes. PTCs partially overlap with super-enhancers, with about 20% of super-enhancers falling within PTCs. However, additional uniquely PTC-associated genes are highly expressed, more so than super-enhancer unique genes (Klonizakis et al. 2023). We have applied SEGCOND to datasets derived from three time points of a CEBPA-induced B-cell to macrophage transdifferentiation system (Rapino et al., 2013). The system represents a highly efficient and homogenous differentiation process, enabling the study of enhancer clusters in a cell-fate conversion context.

Here we set out to investigate how individual enhancers within selected PTCs affect B-cell to macrophage transdifferentiation and the levels of target gene expression. Studying the role of individual enhancers in PTCs associated with *IRF8* and *FOS,* using single and combined enhancer excisions revealed additive, synergistic, and even antagonistic interactions. Our findings also indicate that the enhancer clusters studied behave distinctly in different stages of the immune cell fate conversion, suggesting complex regulatory modes that are likely shaping the differentiation process itself.

## Results

### PTCs dynamically form and dissociate during an immune cell fate switch

To decipher how large enhancer assemblies act during a cell fate conversion process we used a human B-ALL cell line (BLAER) containing an estrogen (E2) inducible CEBPA construct whose activation results in macrophage transdifferentiation after seven days (Rapino et al. 2013)(Fig. 1A). For this, we first identified three-dimensional enhancer clusters at three time points during transdifferentiation (Day 0, Day 1 and Day 7), employing SEGCOND, a previously developed computational pipeline. SEGCOND identified approximately 300 PTCs per differentiation stage, with each PTC consisting of genomic regions of about 200 kilobases in size (Klonizakis et al. 2023).

**Figure 1.**
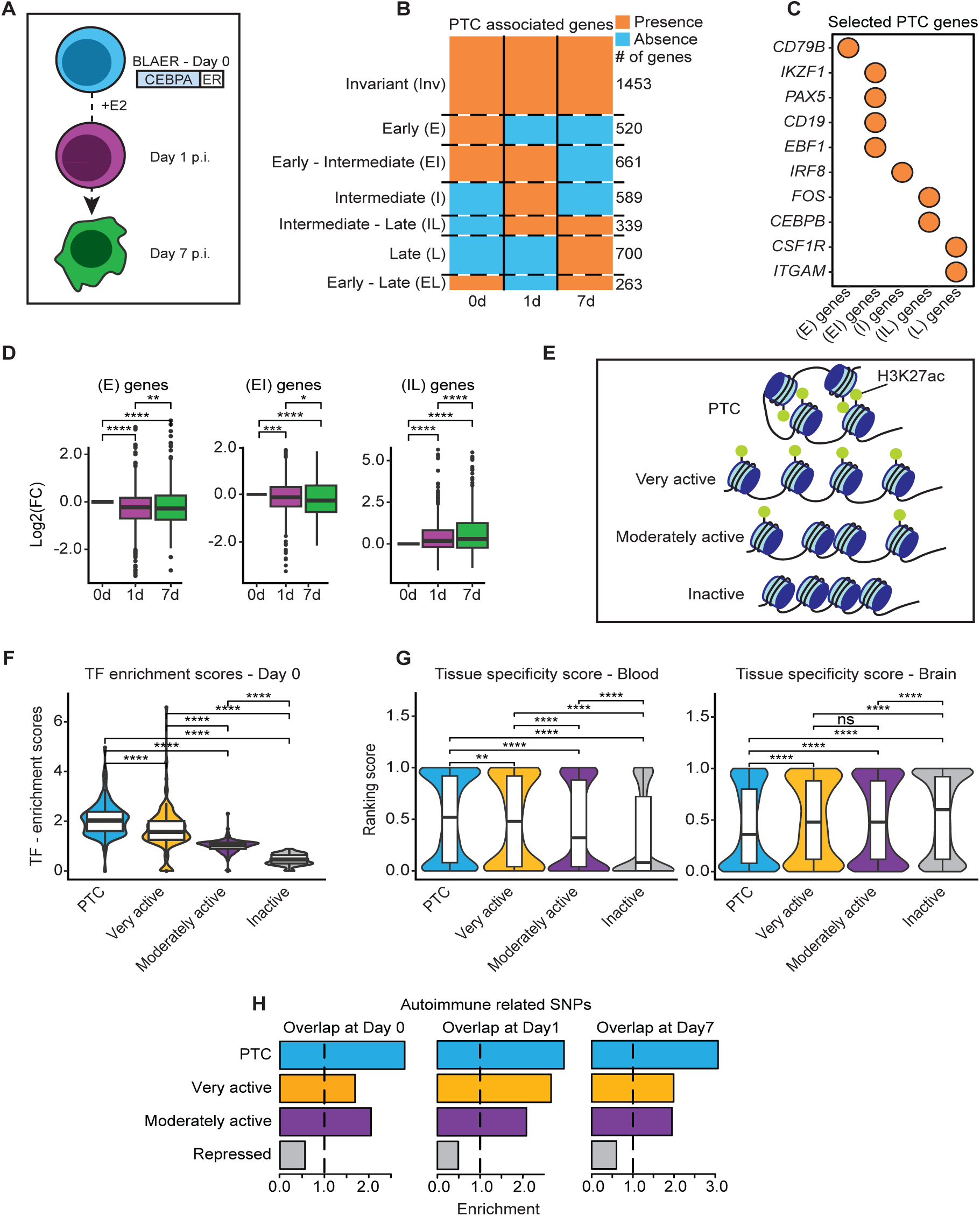
SEGCOND–identified Putative Transcriptional Condensates (PTCs) are dynamic and harbor highly expressed, specialized genes. A. Overview of CEBPA-induced B-cell to macrophage transdifferentiation. BLAER cells, a B-cell ALL cell line derivative, express CEBPA fused with the estrogen receptor (ER). Upon the addition of E2 (b-estradiol), cells transdifferentiate into macrophages over seven days (Rapino et al., 2013). PTCs were previously identified for Day 0, Day 1 and Day 7 cells (Klonizakis et al. 2023). B. Clustering of PTC-associated genes. Expressed protein-coding genes, with an H3K27ac decorated promoter, were clustered according to the timepoint they were associated with a PTC in seven different groups. The PTC clusters were termed “Invariant” (Inv), “Early” (E), “Early-Intermediate” (EI), “Intermediate” (I), “Intermediate - Late” (IL), “Late” (L) and “Early-Late” (EL). C. B-cell and macrophage-specific genes and their association with PTC clusters. Circles indicate gene presence within the corresponding cluster. D. Expression dynamics of PTC-associated genes. DESEq2 variance stabilized counts (Love et al., 2014) of genes grouped in Fig. 1B are depicted for three of the seven clusters. Values of each gene are normalized to their Day 0 expression value. The Wilcoxon signed-rank test was used to determine statistically significant differences (ns p-value > 0.05; * p-value <= 0.05; ** p-value <= 0.01; *** p-value <= 0.001; **** p-value <= 0.0001). E. Separation of the genome into four distinct groups via SEGCOND. Briefly, SEGCOND first partitions the genome into large segments, using ATAC-seq, H3K27ac, H3K4me3, and CEBPA ChIP-seq datasets. Segments are then scored for the presence of H3K27ac decorated, accessible sites (enhancers) with a p-value and a log2FC value. P-values of <= 0.05 indicate enrichment or depletion of enhancer sites compared to random, background genomic regions, while log2FC values indicate whether segments contain more, or less, enhancers than expected by chance. Enhancer-enriched segments (p-value <= 0.05) that interact strongly in three-dimensional space, as determined by Hi-C data, are deemed as “PTC” segments. Enhancer-enriched segments that do not form 3D hubs are termed “Very Active” segments whereas segments that show no enhancer enrichment (p-value > 0.05) but a positive log2FC score are termed “Moderately active” segments. Finally, repressed segments are depleted of enhancer elements (p-value > 0.05 & log2FC <= 0). F. Transcription factor-target enrichment scores for the four SEGCOND genomic groups at Day 0 of transdifferentiation. Transcription factor–target pairs were downloaded by the hTFtarget database (Zhang et al., 2020). An enrichment score, for each transcription factor, was used to determine whether its target genes are over-represented in any of the SEGCOND genomic groups. Statistically significant differences were determined via a Wilcoxon rank-sum test (ns p-value > 0.05; * p-value <= 0.05; ** p-value <= 0.01; *** p-value <= 0.001; **** p-value <= 0.0001). G. Blood and brain tissue-specificity scores of genes associated with SEGCOND genomic categories at Day 0 cells. GTEx data were utilized to assign two scores to each gene, in the 0-1 interval, indicating whether the gene is preferentially expressed in blood or brain tissues respectively (see Method for details). Statistically significant differences were determined via a Wilcoxon rank-sum test (ns p-value > 0.05; * p-value <= 0.05; ** p-value <= 0.01; *** p-value <= 0.001; **** p-value <= 0.0001). H. Overlap enrichment of SEGCOND genomic categories with single nucleotide polymorphisms (SNPs) associated with autoimmune disorders. Enrichment was determined based on a permutation analysis. The dashed line corresponds to the baseline enrichment value of 1, indicating no significant overlap enrichment.

We then analyzed the properties of genes associated with our identified PTCs at the early and late stages of transdifferentiation. We clustered genes into 7 groups according to the timepoint in which they were found to be linked with a PTC (Fig. 1B). Of 4524 genes identified in at least one time point, 1453 genes were found within a PTC throughout the whole process (“Invariant”). The majority of genes (3071) exhibited a dynamic behavior, 520 being associated with a PTC at Day 0 (“Early”), 589 at Day 1 (“Intermediate”), and 700 at Day 7 “Late”. Moreover, 661 genes were found in PTCs at both Day 0 and 1 (“Early-Intermediate”), while 339 were shared in Day 1 and 7 PTCs (“Intermediate-Late”). These data show that the association of genes with PTCs during transdifferentiation is stage-dependent and a dynamic process (Fig. 1B).

We next determined whether the genes associated with PTCs in different clusters correlate with the cells’ differentiation state. At the early stages of transdifferentiation, cells adopt a B-cell identity, which is replaced by a myeloid one as transdifferentiation progresses. Indeed, “Early-Intermediate” membership correlates with B-cell functions while “Intermediate-Late” and “Late” correlate with myeloid functions (Supplementary Fig. 1A). Several B-cell identity genes such as *IKZF1* (Heizmann et al., 2013)*, CD19* (Wang et al., 2012) and *EBF1* (Hagman et al., 2012) are members of Day 0 and Day 1 PTCs, while myeloid related genes such as *IRF8* (Tamura et al., 2000)*, FOS* (Cai et al., 2007)*, CEBPB* (Friedman, 2007) and *CSF1R* (Dai et al., 2002) are found in PTCs at Day 1 and/or Day 7 of transdifferentiation (Fig. 1C). Finally, cell cycle regulation GO terms are overrepresented at different stages, reflecting the fact that BLAER cells become quiescent towards their transdifferentiation to macrophages (Rapino et al., 2013) (Supplementary Fig. 1A).

We have previously shown that PTC-associated genes are highly expressed (Klonizakis et al., 2023). However, it is unclear whether the expression of PTC-associated genes varies over time and whether it is reflected by the kinetics of B-cells to macrophage conversion. We therefore examined expression levels of genes within PTC groups, using their Day 0 values as a reference. This showed that the expression levels of genes associated with PTCs in early stages of transdifferentiation (“Early”, “Early-Intermediate”) decrease after induction, while those of genes associated with PTCs 1 or 7 days after induction (‘Intermediate-Late’ or ‘Late’) increase, as expected (Fig. 1D and Supplementary Fig. 1B).

In conclusion, a large proportion of PTCs exhibit a dynamic behavior during transdifferentiation and are associated with lineage-specific genes and cell cycle genes involved in the cell fate conversion process.

### Genes associated with immune cell PTCs show complex regulation, tissue preference, and disease associations

PTCs harbor a set of lineage-specific, highly expressed genes. We set to further investigate the properties of these genes further and to compare them with other expressed genes in our system that are not under the same multi-enhancer control. To do so, we used SEGCOND to separate the genome into four categories, based on chromatin accessibility (ATAC-seq) and H3K27ac-decorated enhancers (Fig. 1E). “PTCs” are regions identified by SEGCOND (5,5% of genome), “Very Active” regions are enriched in enhancer elements that do not form three dimensional hubs (2,5% of genome), “Moderately Active” are regions neither enriched nor depleted from enhancer elements that also harbor active genes (17,8% of genome) and “Inactive” regions are depleted from enhancer elements and mainly harbor not expressed genes (74,2% of genome) (Fig. 1E, See “Materials and Methods” for more details).

Whether enhancer hubs in the eukaryotic nucleus are established through the association of multiple, diverse transcription factors or a small set of master transcriptional regulators is an open question. To explore these hypotheses, we identified transcription factors (TFs) that are expressed in our cell system and matched them to their target genes, utilizing information from the hTFtarget database (Zhang et al., 2020). We calculated an enrichment score for each SEGCOND-defined genomic category and each transcription factor, quantifying whether target genes of a TF are overrepresented within a genomic category. We found PTCs to be enriched for target genes of multiple distinct TFs, more so than other regions, consistently across transdifferentiation stages (Fig. 1F, Supplementary Fig. 1C). These results indicate that PTC-associated genes are targeted by a higher variety of transcription factors than other active genes.

The complex regulation of PTC-associated genes suggests that they are highly specialized, being mostly active in selected cell types. To address this possibility, we utilized datasets from the GTEx database (Lonsdale et al., 2013) using as controls genes within “Very Active” “Moderately Active” and “Inactive” regions. We also included genes within “Inactive” regions as a negative control. To quantify how specialized the expression pattern of a given gene is, we created a custom “blood” tissue-specificity score. The score ranges from 0 to 1 and describes whether a gene’s expression is higher in the “blood” GTEx samples than in 25 other types of tissues. 1 indicates that a gene’s expression is highest in blood samples while 0 indicates it is at its lowest. Using this score, we observed a significant enrichment for PTCs to contain genes that are preferentially expressed at higher levels in blood tissues, more so than genes in the “Very” and “Moderately Active” regions (Fig. 1G). We then repeated the analysis for “brain” related GTEx samples and observed significant differences between the clusters but in the opposite direction, i.e., the lowest association with PTCs (Fig. 1G). These observations indicate that PTC-associated genes identified in our transdifferentiation system are predominantly active in blood cell types.

The observation that PTCs are associated with a set of specialized genes active in blood tissues raised the possibility that they are linked to blood-related diseases, such as autoimmunity. To this end, we isolated autoimmune-related SNPs from a published study (Maurano et al., 2012) and calculated an enrichment score, using a permutation approach, for all four genomic categories (see Methods). The results show that autoimmune-related SNPs are strongly enriched in PTCs across all time points (Fig. 1H), compared to the other genomic regions that also harbor active genes. To ensure this tendency is specific to autoimmune-related SNPs, we repeated the same analysis using SNPs associated with neurological and behavioral traits and observed no enrichment within PTCs (Supplementary Fig. 1D).

In conclusion, using our immune cell system, SEGCOND identified a set of specialized genomic regions that are under regulatory control of multiple distinct TFs and that are associated with genes involved in autoimmune diseases.

### The transcriptional condensate-associated component BRD4 overlaps with selected PTCs

An interesting question is whether PTCs are associated with transcriptional condensates. BRD4 is a protein component of transcriptional condensates that has been described to co-localize with super-enhancers using DNA-FISH and BRD4 immunofluorescence experiments (Sabari et al., 2018). Therefore, to investigate whether BRD4 also associates with PTCs, we monitored its presence in the *IKZF1* and *IRF8* associated PTCs, as examples representing a B cell and a macrophage PTC, respectively (Heizmann et al., 2013) (Tamura et al., 2000).

We first focused on the 50.14-50.42 Mb PTC of *IKZF1* on chromosome 7, as identified by Hi-C, ATACseq, and H3K27ac ChIP-seq (Fig. 2A). The PTC is active at both Day 0 and Day 1 of transdifferentiation and overlaps with a super-enhancer (Fig. 2A). We marked the *IKZF1* PTC in samples of fixed uninduced BLAER cells with a DNA-FISH probe and then stained the cells with a BRD4 antibody. This showed that they co-localize (Fig. 2B). To evaluate whether the degree of the overlap is more than expected by chance, we developed a computational approach. It first determines whether the DNA-FISH signal correlates with the BRD4 signal on a fixed window around the center of DNA-FISH spots. It then evaluates whether the BRD4 signal is higher in the DNA-FISH spots’ centers than in randomly allocated spots around cell nuclei (see Methods for details). Our analysis revealed that the DNA-FISH signal of uninduced cells correlated significantly with the BRD4 signal (Fig. 2C) and that BRD4 is enriched on DNA-FISH centers when compared with randomly allocated centers (Fig. 2D). Similar results were obtained with Day 1-induced cells after targeting the same locus (Supplementary Fig. 2A-2D).

**Figure 2.**
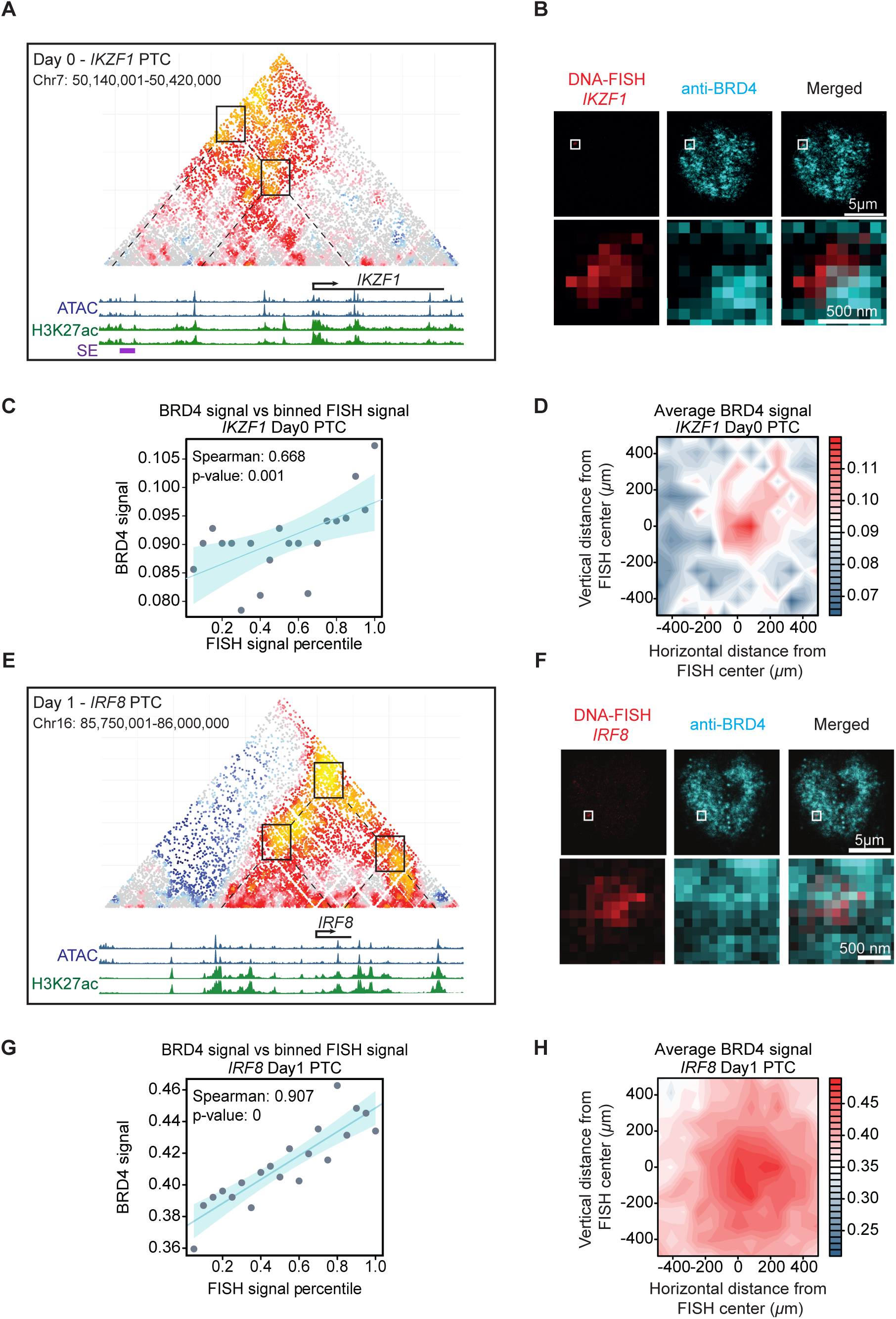
The transcriptional condensate component, BRD4, overlaps with selected PTCs. A. Overview of the *IKZF1* PTC at Day 0 non-induced cells. SHAMAN (Mendelson Cohen et al., 2017) Hi-C normalized values are depicted, with red and yellow points indicating contact enrichment between loci. ATAC-seq and H3K27ac ChIP-seq tracks of two biological replicates are also illustrated. Boxes correspond to H3K27ac decorated elements that interact strongly (dashed lines) in 3D space. A super-enhancer (SE, purple box) within the PTC is also identified by the ROSE algorithm (Whyte et al., 2013). B. DNA-FISH coupled with BRD4 immunofluorescence, targeting the *IKZF1* PTC at Day 0 non-induced cells. BRD4 signal (in cyan) overlaps with the *IKZF1* PTC (in red). The second row of images corresponds to the zoomed region falling within the highlighted area (white frame). C. DNA-FISH and BRD4 signal correlation analysis of Day 0 *IKZF1* PTC data. DNA-FISH pixel intensity values around identified FISH spots were binned in twenty percentiles. Matching BRD4 intensity values from the same pixels were assigned to each bin with the median of all BRD4 values depicted. D. Contour plots of BRD4 signal enrichment over *IKZF1* Day 0 DNA-FISH centers. DNA-FISH spot centers were identified as described in the “Materials and Methods”. A window of 11x11 pixels was centered at each FISH spot and BRD4 intensity values were extracted for each position and each FISH spot. The median BRD4 intensity values, across all DNA-FISH spots, were calculated for each position of the 11x11 window. The process was repeated for randomly allocated FISH centers. Red/blue color codes correspond to the relative enrichment of BRD4 intensity compared to randomized FISH spots. E. Overview of the *IRF8* PTC on Day 1 of transdifferentiation. Same as in Fig. 2A. ROSE doesn’t detect a super-enhancer within the PTC. F. DNA-FISH coupled with BRD4 immunofluorescence, targeting the *IRF8* PTC at Day 1 cells. Same as in Fig. 2B. G. DNA-FISH and BRD4 signal correlation analysis for Day1 *IRF8* PTC data. Same as in Fig.2C. H. Contour plots of BRD4 signal enrichment over *IRF8* Day 1 DNA-FISH centers. Same as in Fig.2D.

We next studied the *IRF8* PTC on chromosome 16 which is only active at Day 1 of transdifferentiation (Fig. 2E). Although *Irf8* super-enhancers have been reported in murine cells (Liu et al., 2023a), the ROSE algorithm didn’t identify one within the *IRF8* PTC in our system (Fig. 2E). We first evaluated whether BRD4 hubs assemble on this region predicted by SEGCOND, and observed that they indeed overlap with the *IRF8* PTC in Day 1-induced cells (Fig. 2F). Applying our computational analysis pipeline to DNA-FISH intensity and BRD4 signal at this locus yielded a highly significant correlation at Day 1 of transdifferentiation (Fig. 2G), as well as an enrichment of BRD4 signals in *IRF8* DNA-FISH centers compared to randomly allocated spots (Fig. 2H). No such findings were made with Day 0 cells, as the PTC is not active at that stage (Supplementary Fig. 2E & 2F).

In conclusion, our results indicate that BRD4 hubs assemble at PTCs that are specifically formed at either Day 0 and 1 (*IKZF1*) or Day 1 cells (*IRF8*), in line with the idea that they correspond to enhancer regions that form condensates

### Two enhancers within the PTC of *IRF8* confer robustness of transdifferentiation and high expression levels of IRF8

To study how enhancers within PTCs modulate gene expression and transdifferentiation, we chose the *IRF8* PTC as a test case. *IRF8* is a known myeloid regulator (Tamura et al., 2000) which is upregulated approximately 16-fold in our transdifferentiation system at Day 1 post-CEBPA induction. The *IRF8* PTC contains multiple enhancers, of which two, located at -69kb and +83kb from the *IRF8* start site, are bound by CEBPA and its obligate partner PU.1 (Van Oevelen et al., 2015)(Heinz et al., 2010)(Fig. 3A). The two individual enhancers show an increase of their chromatin accessibility and H3K27ac decoration at Day 1 compared to uninduced cells. They thus prove ideal candidates for excision as their deletion should not cause cell lethality in uninduced cells and as they are not bound by CTCF, eliminating a possible confounding parameter due to it having a topological role (Ong & Corces, 2014) (Figure 3A).

**Figure 3.**
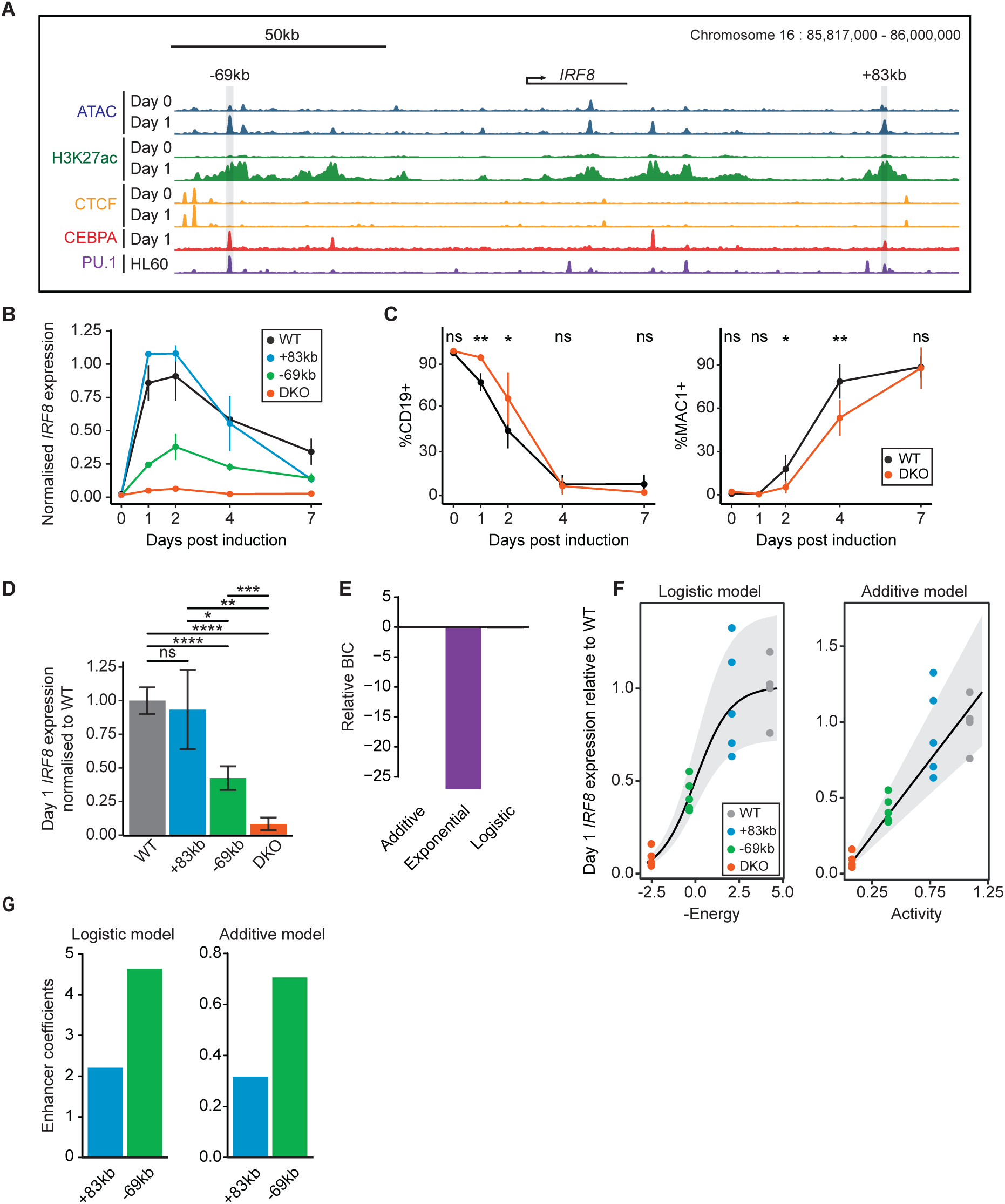
The -69kb and +83kb *IRF8* enhancers provide robustness to *IRF8* expression levels and transdifferentiation. A. Overview of the *IRF8* PTC chromatin landscape at Day 0 and Day 1 cells. ATAC-seq, H3K27ac, and CTCF ChIP-seq tracks of Day 0 and Day 1 cells are depicted. Two enhancers, -69kb and +83kb away from the *IRF8* transcription start site were chosen for excision, as they show an increase in H3K27ac decoration and chromatin accessibility at Day 1 cells, are not bound by CTCF and are bound by C/EBPa in Day 1 cells and PU.1 in HL60 cells. B. *IRF8* expression kinetics during transdifferentiation. *IRF8* expression levels were monitored via RT-qPCR and were normalized against *GUSB* expression levels for each timepoint. C. Transdifferentiation kinetics of *IRF8* DKO cells compared to BLAER cells, monitored via FACS. CD19+ and MAC1+ cells were evaluated across 5 timepoints. The mean ± s.d is depicted for each timepoint. Statistical significance was determined using a paired Student’s t-test (ns p-value > 0.05; * p-value <= 0.05; ** p-value <= 0.01; *** p-value <= 0.001; **** p-value <= 0.0001). D. Day 1 *IRF8* expression levels of all generated KO lines and WT cells. *IRF8* expression levels were determined via RT-qPCR and normalized against *GUSB* expression levels. Consequently, expression values from each transdifferentiation batch were normalized to the average WT expression values of the same batch. Statistical significance was determined using a Student’s t-test (ns p-value > 0.05; * p-value <= 0.05; ** p-value <= 0.01; *** p-value <= 0.001; **** p-value <= 0.0001). E. Evaluation of best statistical model fit for Day 1 *IRF8* expression data. The Bayesian Information Criterion (BIC) was used to determine which model better fits the expression data. BIC values of the additive loci were used as a reference point. A BIC difference greater than 2 indicates a better fit. Both the additive and logistic models were found to be valid. F. Logistic and additive model fits to *IRF8* Day 1 expression data. Black lines denote the models’ average predicted expression values for each enhancer combination, with the shaded area corresponding to 0.1-0.9 quantiles. Individual expression values, normalized to WT values, are depicted as points. The x-axis of the logistic model corresponds to the energy being conferred towards increasing transcriptional output by the addition of each enhancer. The x-axis of the additive model corresponds to the activity of each enhancer which is proportional to gene expression. G. Logistic and additive model enhancer coefficients for *IRF8* Day 1 expression data. The enhancer coefficients determine enhancer potency towards increasing expression levels. Both models showcase that the -69kb enhancer is stronger than the +83kb enhancer at this transdifferentiation stage.

We proceeded in excising the -69kb and +83kb enhancers by CRISPR separately and in combination, producing three knockout (KO) lines (Fig. 3A). These lines were induced to transdifferentiate and monitored at five time points for the expression of *IRF8* via RT-qPCR. We expected that single enhancer excisions would result in a modest reduction of *IRF8* upregulation and that their combined excision would lead to a more severe reduction. However, surprisingly, while deletion of the +83kb enhancer led to no obvious differences in *IRF8* levels at Day 1,2 and 4 of transdifferentiation, excision of the -69kb enhancer led to a strong *IRF8* RNA reduction and their combination to an even more severe decrease at all timepoints after induction (Fig. 3B).

We next measured the protein levels of CD19 and MAC1 (CD11B) during transdifferentiation, permitting us to simultaneously assess the silencing of the B cell program and activation of the macrophage program (Supplementary Fig. 3A). The results obtained showed that excision of the -69kb enhancer slightly but significantly delayed CD19 downregulation at days 1 and 2 p.i. without affecting MAC1 activation (Supplementary Fig. 3B). Excising the +83kb enhancer likewise delayed B cell silencing without affecting the macrophage program (Supplementary Fig. 3B). Strikingly, however, the double knockout (DKO) cells showed a significant delay not only of CD19 silencing but also of MAC1 activation (Fig. 3C). Of note, all three knockout cell lines eventually converted into induced macrophages at Day 7 p.i., as evidenced by their marker expression.

In an attempt to dissect the possible interactions between the -69kb and +83kb enhancers in the *IRF8* PTC, we utilized three previously described statistical models, termed additive, exponential, and logistic (Dukler et al., 2016). Each model assumes a different relationship between the two enhancers, leading to different transcriptional outputs. According to the additive model, each enhancer independently and additively contributes to the strength of the enhancer cluster, while the exponential model suggests that enhancers within a cluster interact, amplifying or dampening each other’s activity. Finally, the logistic model proposes that each enhancer contributes independently to the reduction of an energy threshold important for transcription (Supplementary Fig. 3C, see Methods for more details). We fit the three models in our Day 1 and Day 7 expression data after performing additional transdifferentiation experiments, including two independent -69kb and DKO cell clones. We observed that on Day 1 p.i. *IRF8* levels do not differ significantly between the wild type and the +83kb *IRF8* KO cells, are reduced in -69kb cells and further reduced in DKO cells (Fig. 3D). Both the logistic and additive models were found to be good fits for our Day 1 dataset, as determined by the Bayesian Information Criterion (BIC) (Fig. 3E). The logistic model predicts a sigmoid-like behavior for *IRF8* expression, where even small differences in enhancer activity can lead to sharp reductions in expression levels, while the additive model suggests that the two enhancers’ activity is added up linearly (Fig. 3F). Notably, both models highlight a large difference between the activity of the two enhancers, with the -69kb enhancer being approximately 2.5 times stronger than the +83kb enhancer at Day 1 p.i. (Fig. 3G). At Day 7 p.i. the deletion of either enhancer leads to a reduction in expression, with their combined excision introducing an even stronger decrease (Supplementary Fig. 3D). The best fit was the exponential model, which indicates synergy between the two enhancers (Supplementary Figures 3E & 3F). The model also suggests that the strength of the two enhancers was similar at Day 7 p.i. cells (Supplementary Fig. 3G).

Overall, the two enhancers studied within the PTC of *IRF8* were found to control the gene’s expression differently at distinct stages of transdifferentiation and to cooperate in driving high levels of *IRF8* expression. These effects translate into more complex outcomes at the level of transdifferentiation kinetics, suggesting thresholds of IRF8 important for both the silencing and activation of lineage markers.

### Two enhancers in the PTC of *FOS* show time-dependent negative and positive interactions during induced transdifferentiation

To determine whether the observed behavior of the two *IRF8* enhancers is a more general property of PTCs we analyzed the *FOS* locus, which is found within a PTC that forms at Days 1 and 7 of transdifferentiation and that is also captured by the super-enhancer algorithm (Figure 4A). FOS has been reported to play a role in monocyte specification through dimerization with CEBPA (Cai et al., 2007). Inspection of the *FOS* PTC revealed a -3.3kb and a +8.9kb enhancer bound by CEBPA at 1 Day p.i. but not by CTCF (Fig. 4A). These enhancers were again excised in the BLAER cell line individually or in combination, generating three derivative lines.

**Figure 4.**
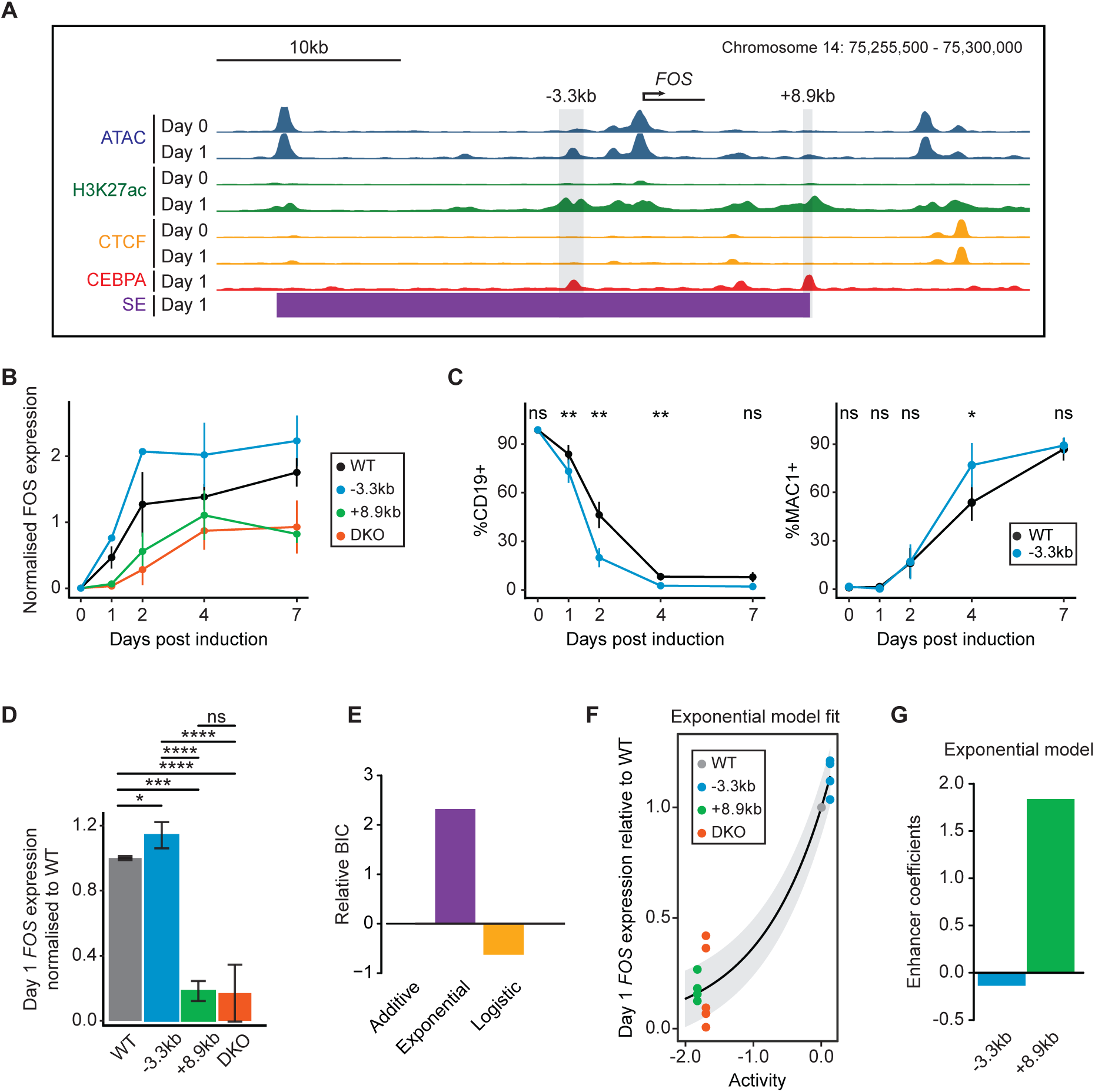
The -3.3kb and +8.9kb *FOS* enhancers antagonize at Day 1 cells, delaying transdifferentiation kinetics. A. Overview of the *FOS* PTC at Day 0 and Day 1 cells. ATAC-seq, H3K27ac, and CTCF ChIP-seq tracks of Day 0 and Day 1 cells are shown. Two enhancers, -3.3kb and +8.9kb from the *FOS* transcription start site were chosen for excision, as they become more accessible, show an increase in H3K27ac signal at Day 1 cells, are not occupied by CTCF in any timepoint and are occupied by C/EBPa in Day 1 cells. A super-enhancer called by the ROSE algorithm is also identified in the *FOS* PTC, containing the two enhancer elements. B. *FOS* expression levels throughout transdifferentiation. *FOS* expression levels were measured via qPCR and were normalized against *GUSB* expression levels C. Transdifferentiation kinetics of *FOS* -3.3kb KO compared to BLAER cells, assessed via FACS. The mean ± s.d of CD19+ and MAC1+ cells is depicted for each timepoint. Statistical significance was determined using a paired Student’s t-test (ns p-value > 0.05; * p-value <= 0.05; ** p-value <= 0.01; *** p-value <= 0.001; **** p-value <= 0.0001). D. Day 1 *FOS* expression levels of KO and WT cells. *FOS* expression levels were measured via RT-qPCR and normalized to *GUSB* expression levels. Consequently, expression values from each transdifferentiation batch were normalized to the average WT expression values of the same batch. Statistical significance was determined using a Student’s t-test (ns p-value > 0.05; * p-value <= 0.05; ** p-value <= 0.01; *** p-value <= 0.001; **** p-value <= 0.0001). E. Selection of best model fit for Day 1 *FOS* expression data. As in Fig. 3E. The exponential model was the better fit to the data. F. Exponential model fit to *FOS* Day 1 expression data. Same as in Fig. 3F. G. Exponential model enhancer coefficients for *FOS* Day 1 expression data. The -3.3kb enhancer is assigned a negative coefficient as it dampens the activity of the +8.9kb enhancer.

We next measured expression levels of *FOS* across the five time points of transdifferentiation (Fig. 4B). Unexpectedly, deletion of the -3.3kb enhancer led to a transient up-regulation of *FOS* at Days 1 and 2 p.i., suggesting that at these timepoints the -3.3kb enhancer functions as a repressor. This repression is not mediated by the polycomb complex as the -3.3kb enhancer was not found to be decorated with the repressive histone mark H3K27me3 (Supplementary Fig. 4B). In contrast, excision of the +8.9kb enhancer led to a pronounced reduction in *FOS* expression at Days 1 and 2 p.i, as in DKO cells.

We then monitored the transdifferentiation dynamics of the cell line derivatives by FACS. Surprisingly, excision of the -3.3kb enhancer substantially and significantly accelerated both the silencing of CD19 and activation of MAC1 (Fig. 4B). In contrast, ablation of the +8.9kb enhancer did not alter the transdifferentiation kinetics (Supplementary Fig. 4A) while its deletion in DKO cells counteracted the observed acceleration caused by the -3.3kb enhancer excision (Supplementary Fig. 4A). These findings suggest that when *FOS* expression levels range between those observed in +8.9kb/DKO cells and WT cells, transdifferentiation can proceed normally. In contrast, elevated levels of FOS, exceeding those in WT cells, accelerate the kinetics of transdifferentiation.

We generated a second set of cell clones for the three conditions, to exclude any potential clone-specific biases. Once again, we observed an increase of *FOS* expression at Day 1 p.i. in the -3.3kb clones and a sharp decrease in the +8.9kb and DKO clones (Fig. 4D). We then applied statistical modeling to our Day 1 expression data and determined that the exponential model results in the best fit (Fig 4E). Under the assumption of this model, the two enhancer elements act synergistically to modulate each other’s activity (Fig. 4F). However, in this case, the -3.3kb enhancer displays a negative coefficient (Fig. 4G) as it antagonizes the activity of the +8.9kb enhancer. Repeating the same analysis for Day 7 cells showed that the DKO cells exhibited a decrease of *FOS* expression but not the individual KO cell lines (Supplementary Fig. 4C). In contrast to Day 1 data, both the logistic and additive models were found to fit the Day 7 expression data better (Supplementary Figures 4D & 4E). Under the latter models, the -3.3kb enhancer now functions as an activator rather than as a repressor, although weaker than the +8.9kb enhancer (Supplementary Fig. 4F). This result suggests that there is a switch in the behavior of the *FOS* PTC between Day 1 and 7 of transdifferentiation.

In conclusion, our data show that the -3.3kb FOS enhancer antagonizes the activity of the +8.9kb enhancer on Day 1, dampening the transdifferentiation kinetics and FOS expression. The antagonism is Day 1 stage-specific, with the two enhancers cooperating in Day 7 induced cells and providing robustness to *FOS* expression levels.

## Discussion

Here we studied putative transcriptional condensates (PTCs) identified by SEGCOND (Klonizakis et al., 2023) during an induced cell fate conversion. Unlike other *in silico* methods, SEGCOND first partitions the genome into large segments and then identifies PTCs through the integration of Hi-C data. Our pipeline revealed enhancer elements far apart across a given chromosome and interacting to form large clusters of 3D hubs, regulating gene expression. PTCs were found to be enriched for lineage instructive genes, as has been described for other large enhancer clusters, including LCRs and super-enhancers (Grosveld et al. 1987) (Whyte et al. 2013) (Parker et al. 2013) (Madsen et al. 2020). More specifically, our results showed that individual enhancers within the PTCs of *IRF8* and *FOS*, genes encoding TFs implicated in regulating immune cell development, interact in differentiation stage-dependent complex manners.

PTCs identified in our immune cell transdifferentiation system are associated with highly specialized genes, preferentially expressed in blood cells. Moreover, they are enriched for SNPs that have been linked to autoimmune disorders. For example, the *IRF8* +83kb enhancer described here contains an SNP, termed rs9927316, associated with rheumatoid arthritis (Freudenberg et al., 2015). As the *IRF8* PTC is not detected by other methods, like the ROSE algorithm, SEGCOND could represent an attractive framework for the discovery of cell type-specific disease variants. Moreover, SEGCOND allows the inspection of single nucleotide polymorphisms in a three-dimensional context. This could reveal previously overlooked links between SNPs within regulatory genomic regions and target genes that may be distant in linear space.

Since BRD4 is known to associate with super-enhancers and to form condensates, (Sabari et al. 2018) we used its co-localization with PTCs as a proxy for their participation in transcriptional condensates. Indeed, BRD4 could be observed to overlap with the *IKZF1* PTC, which is also called a super-enhancer by the ROSE algorithm. However, we detected BRD4 at the *IRF8* PTC in our transdifferentiation model despite it not qualifying as a super-enhancer. Our results suggest that the ROSE algorithm underestimates the number of genes controlled by BRD4 hubs in our cell system. Whether PTCs associate with condensates in live cells could not be determined since DNA-FISH requires cell fixation. Nevertheless, future experiments resolving this question appear feasible, since MED1, another component of transcriptional condensates, was recently shown to be important for the frequency of transcriptional bursts in mouse embryonic stem cells (Du et al., 2024). Using live-cell microscopy, it was concluded that MED1-containing condensates associate with the *Sox2* locus and that their proximity correlates with the transcriptional activity of the gene (Du et al., 2024). Therefore, at this stage, SEGCOND can only pinpoint candidate gene loci that support the formation of BRD4 assemblies into transcriptional condensates.

Our knockout data showed that although both the -69kb and +83kb *IRF8* enhancers control *IRF8* expression at Day 1 of transdifferentiation, the -69kb enhancer is about 2.5 fold stronger than the +83kb enhancer. This implies that the activity of the -69kb enhancer overshadows the activity of the +83kb enhancer, “saturating” *IRF8* expression levels. However, in the absence of the -69kb enhancer the +83kb enhancer remains active, representing a mechanism that safeguards *IRF8* expression (Figure 5A). Therefore, these findings show that the *IRF8* PTC confers robustness to the gene’s expression and transdifferentiation, as only the double enhancer excision substantially impaired the two parameters. Such expression thresholds of target genes may also be critical for other differentiation processes, with enhancer clusters functioning as safeguards that ensure expression levels above a critical threshold. Similar findings have been described for distinct enhancer pairs that determine the expression of *Gli3* and *Shox2,* factors required for murine limb development. Here, while excision of single enhancers of the two genes showed no effect, their combined deletion caused developmental defects (Osterwalder et al., 2018).

**Figure 5.**
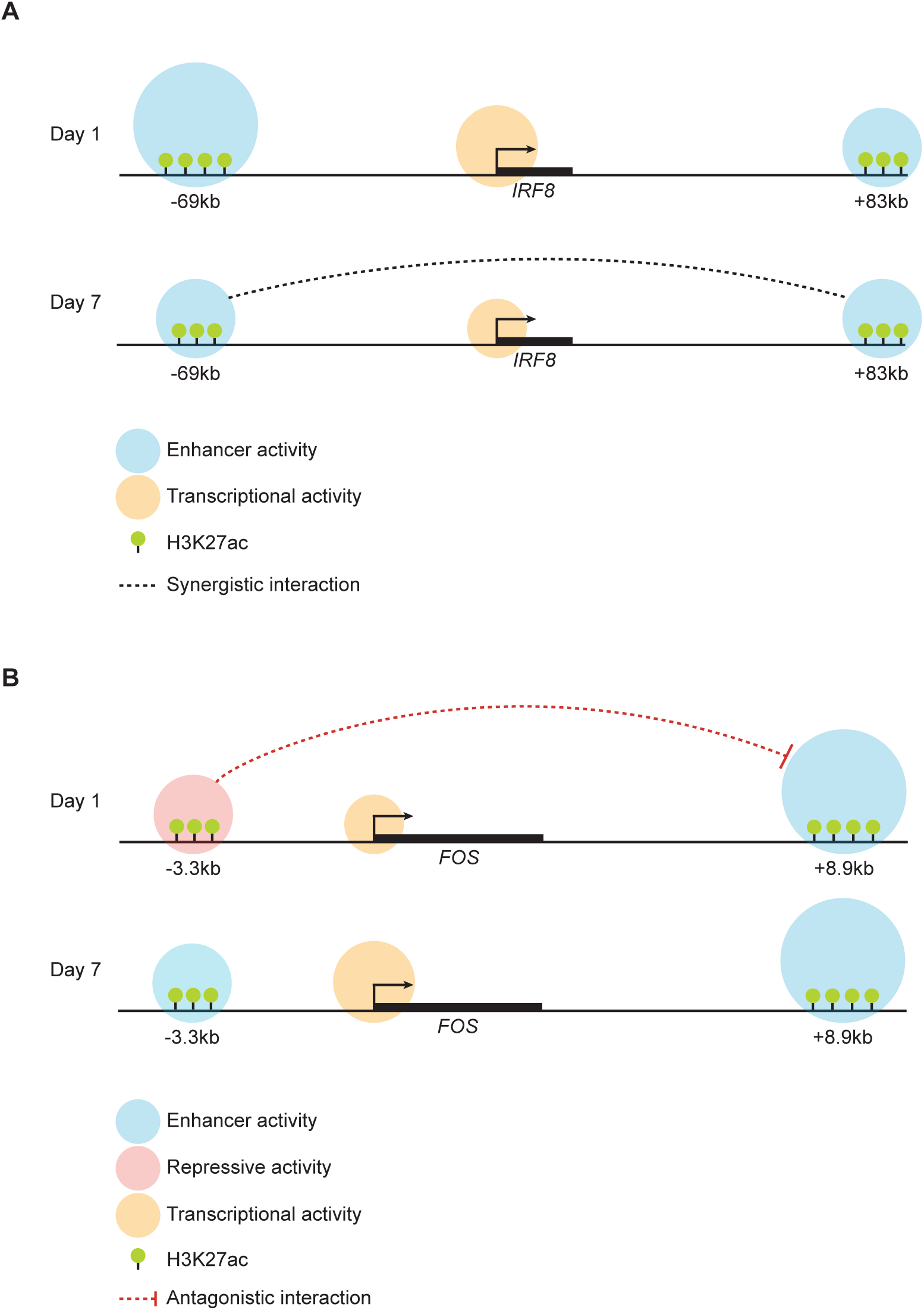
Differentiation stage-restricted activities of enhancer pairs during immune cell transdifferentiation. A. Graphical model for *IRF8* enhancer activity during transdifferentiation. At Day 1 cells, the -69kb enhancer is stronger than the +83kb enhancer, leading to high *IRF8* expression levels and a “masking” of +83kb enhancer activity. However, the presence of both enhancers forms a safeguard mechanism, ensuring that *IRF8* expression levels are maintained above a threshold needed for efficient transdifferentiation. At Day 7 cells the two enhancers display similar activity levels and synergize, driving higher *IRF8* expression levels than expected under an additive model. B. Graphical model for *FOS* enhancer activity during transdifferentiation. At Day 1 cells the -3.3kb enhancer antagonizes the activity of the +8.9kb enhancer, leading to a decrease in *FOS* expression and a concomitant decrease of transdifferentiation kinetics. This behavior is switched at Day 7 cells, where the two enhancers cooperate independently, driving high *FOS* expression levels.

The functional characterization of individual enhancers of the FOS PTC yielded surprisingly different results. Hence, instead of a synergism between its -3.3kb and +8.9kb enhancers, it revealed an *antagonistic* relationship in Day 1 induced cells (Figure 5B). A possible explanation for the observed antagonism is that the -3.3kb element functions as a silencer on Day 1 of transdifferentiation. However, the -3.3kb enhancer is decorated by the activating mark H3K27ac but not with the repressive H3K27me3 mark at this stage. An alternative explanation is that the two enhancers locally recruit large amounts of transcription factors that then become repressive. In line with this idea is the observation that the formation of transcriptional condensates by specific synthetic transcription factors can inhibit transcription (Trojanowski et al. 2022). Such a mechanism could be relevant for the calibration of gene expression levels. Thus, limiting the maximum rate of mRNA production could lock the expression level of a gene in a concentration range required during a particular stage of differentiation. As such, in our model, the excision of the -3.3kb enhancer led to higher FOS levels and the acceleration of transdifferentiation.

Our results suggest that enhancer clusters can act as transcription factor “control panels”, leading to different transcriptional outputs depending on the availability of stage-specific transcription factors and/or co-factors. Accordingly, the relationship between individual enhancers of each cluster proved to be dynamic, with the two *IRF8* enhancers synergizing in Day 7 induced cells (corresponding to mature macrophages), but not in Day 1 induced cells (likely corresponding to early myeloid precursors), where they independently add up their activity levels. In contrast, while the two *FOS* enhancers also cooperate in Day 7-induced cells, they antagonize each other in Day 1-induced cells. A similar stage-dependent activity has also been reported for the murine *Irf8* enhancer cluster, where Individual enhancers control pDC1 differentiation at distinct stages (Durai et al., 2019) and also synergize during a short developmental window (Liu et al., 2023b). In summary, applying SEGCOND to an immune transdifferentiation system has helped us to discover novel enhancer clusters (PTCs) missed by other algorithms. Our discovery that individual enhancers within PTCs not only act synergistically but also antagonistically may have uncovered an as-yet-overlooked new principle of enhancer interactions.

## Materials and methods

### Application of SEGCOND and datasets used

SEGCOND was applied to datasets derived from an established B-cell to macrophage transdifferentiation system (Rapino et al., 2013), as described in (Klonizakis et al., 2023). H3K27ac, H3K4me3, and H3K27me3 ChIP-seq datasets used throughout the study were downloaded from (Borsari et al., 2020). ATAC-seq, RNA-seq, CTCF ChIP-seq, and Hi-C datasets are derived from (Stik et al., 2020), while CEBPA ChIP-seq datasets are from (Choi et al., 2021). PU.1 ChIP-seq data derived from the HL60 cell line was downloaded from GEO (GSM1010843).

### Clustering of genes within PTCs and functional enrichment analysis

To determine which genes are members of PTCs highlighted by SEGCOND, we employed a gene filtering strategy. We only included protein-coding genes that had an average TPM value of > 1 across the three time points tested. We further asked that genes in enhancer hubs harbored at minimum an H3K27ac peak within a 1000 base pair window from their promoter. Our analysis yielded 4525 genes across all three studied time points. We further separated them into categories according to their time-related PTC membership. GO term analysis was performed in the identified groups by gprofiler2 (Kolberg et al., 2020).

### B-cell and myeloid signature score

Modules of B-cell and myeloid-specific genes were downloaded from a published study (Monaco et al., 2019). We conducted a hypergeometric test in R to determine statistically significant overlaps between genes in PTC association clusters and the isolated modules.

### Separation of genomic categories by SEGCOND

In order to compare the functional characteristics of PTC regions with other meaningful genomic regions, we split the genome into four categories, using SEGCOND. SEGCOND separates the genome into segments and then calculates a log2FC enrichment score and a p-value for each segment based on a user-provided characteristic. In our case, the characteristic of interest was the presence of enhancers. Using SEGCOND’s quantification schemes for enhancer enrichment within each segment, we separated the genome into four distinct region classes as shown below:

“Inactive”: Genomic regions that had a p-value of >0.05 and a log2FC < 0

“Moderately active”: Genomic regions with a p-value >0.05 and a log2FC >= 0

“Very active”: Genomic regions with <=0.05 p-value and a log2FC >= 0

“PTC”: Genomic regions with a <=0.05 p-value, log2FC >= 0 and high Hi-C interaction scores as assessed by SHAMAN (Mendelson Cohen et al., 2017)

### TF-target enrichment scores

To determine whether blood tissue-related transcription factors tend to have more gene targets within PTCs, we utilized the hTFtarget database (Zhang et al., 2020). Transcription factor and target pairs were downloaded and we filtered the table to only pairs known to occur in blood-related tissues. We further filtered the list for transcription factors expressed in our system. We included all protein-coding gene targets, irrespectively from their expression levels, as otherwise we would bias our results towards regions that harbor more active genes. For each transcription factor, we calculated the percentage of total genes regulated by the factor. The “Expected” score of target genes for each TF and each genomic category was thus:

Expected genes = (Percentage of gene targets) x (Number of genes found within a genomic category)

The final enrichment score per genomic category was calculated as:

Enrichment Score = Observed genes / Expected genes

### GTEx tissue specificity scores

GTEx TPM expression data were downloaded (Lonsdale et al., 2013). For each tissue, genes were ranked according to their expression profile. Expression values were substituted by each gene’s rank. In case a tissue had multiple distinct datasets assigned to it (e.g., different brain regions, UV/non-UV exposed skin), gene ranks were averaged. To determine whether a gene is preferentially expressed in Blood tissues a custom score was generated:

The tissue-specific score for gene I (Blood) = # Tissues with a lower Rank than the Blood rank of gene i / # Tissues

The range of values of the above score range from 0 to 1. A value of 1 indicates that a gene’s rank is the highest in blood tissue and shows biased expression towards it. On the contrary, a value of 0 indicates that a gene’s rank is the lowest in blood tissues. The same strategy was applied to get the “Brain” tissue specificity score.

### Enrichment of auto-immune SNPs

Genomic coordinates of SNPs were downloaded from (Maurano et al., 2012) and only autoimmune-related SNPs were kept for further analysis. To determine enrichments of SNPs within different genomic regions, we employed a permutation approach. Using bedtools shuffle (Quinlan & Hall, 2010) coordinates corresponding to the four genomic categories under study were randomly shuffled across the genome a hundred times. After each permutation, overlap with SNPs was calculated and stored. The final enrichment score for each category was:

Enrichment = # Observed SNP Overlap / Average number of SNP overlap with permuted regions

### Preparation of DNA-FISH probes

To target the *IKZF1* and *IRF8* PTCs, in-house fluorescent DNA probes were prepared. BAC clones containing the target loci were ordered (BACPAC genomics, RP11-663L2 and RP11-478M13) and a Nick translation reaction (Roche, Cat 10976776001) was performed where Cy5-labeled UTP nucleotides (Jena, Cat NU-803-CY5-L) were incorporated. The ratio of UTP to non-labeled TTP nucleotides was 30:70. Nick translation was performed following the manufacturer’s protocol. After stopping the nick translation reaction, a small quantity of probes was run on a 1% agarose gel, to evaluate the size of produced DNA fragments. Probes that exhibited a smear of 100-500 base pairs were purified using the Zymo DNA Clean and Concentrator Kit (Zymo Research, Cat D4013) and were used for DNA-FISH experiments.

### DNA FISH coupled with BRD4 immunofluorescence

DNA-FISH was performed as described in (Bolland et al., 2013) with minor modifications. Washes were performed in 24 well plates (Corning, Cat 353047) instead of Coplin jars. Poly-L lysine-coated coverslips were used (Electron Microscopy Sciences, Cat #72292-01) and 200.000 cells were seeded per coverslip. Nitrogen snap-freeze cycles were substituted with dry-ice freeze cycles, with coverslips placed for 15-30 seconds on top of dry ice per cycle. 50ng of in-house labeled probes were used per reaction. To perform BRD4 staining using antibodies, the original protocol was extended. Coverslips were re-calibrated in PBST (PBS with 0,2% Tween 20 Merck, Cat 9005-64-5) for 5 minutes and 50ul of primary antibody (rabbit anti-BRD4, ref: ab128874, 1:50 dilution) in blocking solution (PBST + 4% BSA) was subsequently placed on top of coverslips. Samples were incubated O/N at 4C. The following day, 3 PBST washes of 5 minutes each were performed followed by incubation of 250 ul secondary antibody solution (Alexa Fluor 546, goat anti-rabbit, Cat A11071, 1:500 dilution in PBST + 4% BSA) for 1 hour at room temperature. 3 rounds of PBST washes were repeated, following mounting with a drop of Fluoroshield with DAPI (Sigma, F6057-20ml). Samples were visualized using an SP5 Inverted Leica confocal microscope.

### Computational analysis of images from DNA-FISH & BRD4 IF experiments

Images were processed in R using the EBImage package (Pau et al., 2010). Confocal image stacks of 1024x1024 pixels were processed in the following order:

1. Identification of nuclei. A smoothing Gaussian brush was passed at each DAPI stack (brush size: 31, sigma of 5). Stacks were otsu thresholded and the signal was binarized to identify nuclei. Holes were filled with the fillHull() command.

2. Identification of FISH spots. A smoothing Gaussian brush was passed at each FISH channel stack (brush size: 9, sigma of 5). A stringent thresholding cutoff was picked manually for each set of stacks and was applied to determine DNA-FISH spots. Any spot that didn’t fall within a nucleus was excluded from the analysis. Additionally, nuclei that didn’t contain any FISH spots weren’t considered for downstream analyses.

3. Identification of FISH spot centers. The computeFeatures.moment() command was used to extract centers of identified FISH spots per stack, based on the shape of FISH spots. Centers were then shifted to get the brightest pixel as the central spot. An identifier was assigned to each FISH spot, to track the same FISH spot spanning multiple stacks and only count it once when calculating BRD4 signal overlaps.

4. 3D reconstruction of nuclei and random allocation of spots. Nuclei that harbored FISH spots were stitched in 3D across stacks to reconstruct their total nuclear volume. 50 centers were randomly assigned at each nucleus to determine the background BRD4 signal.

5. Calculation of BRD4 occupancy around FISH centers and random spots. A window of 11x11 was centered across each identified FISH center and stack. Values from the BRD4 channel were extracted and assigned to each entry of the 11x11 matrix. Every time a value is assigned to the 11x11 window, a check takes place to determine whether the window is within a nuclear volume. In the opposite case, a value of 0 is assigned alongside a flag that excludes these entries from further calculations. Finally, an average matrix is calculated for every unique FISH spot spanning multiple image stacks. The same process was repeated for the randomly allocated centers.

6. Generation of contour plots. Median values across all matrix entries of genuine DNA-FISH spots were extracted and plotted as a contour plot with the filled.contour() command. A similar strategy was employed for the randomly allocated spots. The lowest color scale value was set to blue to match the values of the randomly allocated centers. Consequently, values that aren’t blue indicate an enrichment of BRD4 intensity in relation to background spots.

7. Calculation of correlation coefficients. In a later step, we wanted to establish whether the BRD4 signal correlates with the increasing intensity of the DNA-FISH signal. To do so, all BRD4 values within the centered 11x11 windows were extracted and matched to the corresponding FISH-intensity values from the FISH channels. FISH values were binned in 5% percentiles and BRD4 values were assigned to each matching bin. The median values of all BRD4 data points per bin were extracted and plotted.

### Cell culture and transdifferentiation experiments

BLAER cells and enhancer KO derivatives were cultured in RPMI medium (Gibco, catalog no. 22400089) supplemented with 10% fetal bovine serum (Gibco, catalog no. 10100147), 1% glutamine (Gibco, catalog no. 25030081), 1% penicillin/streptomycin mix (Thermo Fisher Scientific, catalog no. 15140122) and 50 µM β-mercaptoethanol (Gibco, catalog no. 31350010). To transdifferentiate BLAER cells and derivatives, 250.000 cells were seeded in 12-well plates in 1mL of growth medium supplemented by 100 nM E2 (b-estradiol), hIL-3, and hCSF-1 (10 ng ml−1). Cells were tested every month for mycoplasma infection and were always found negative.

### Generation of enhancer knock-out cell lines

To generate enhancer KO cell lines, we cloned the px330_mCherry and px330_TagBFP plasmids by inserting mCherry and TagBFP sequences (that lack a BbsI cut site), into a base px330 plasmid (Addgene #98750). The base px330 plasmid was a gift from Jinsong Li (Addgene plasmid # 98750; http://n2t.net/addgene:98750; RRID: Addgene_98750). The final plasmids encode the CAS9 enzyme and the corresponding fluorescent marker while they contain a cloning site for an sgRNA of choice.

To excise an enhancer, we simultaneously targeted two sites flanking it. We thus cloned the following sgRNA pairs (Integrated DNA Technologies), after annealing them, in our px330 vectors, generating a set of two px330 plasmids per enhancer excision:

*IRF8* +83kb, chromosome 16:85,982,380 - 85,982,912 (hg38)

cacc**GCAAAGACAATGAGAAGCGG**, aaac**CCGCTTCTCATTGTCTTTGC** (mCherry)

cacc**GTGTGGCCTCTCGTGTCAGT**, aaac**ACTGACACGAGAGGCCACAC** (TagBFP)

*IRF8* -69kb, chromosome 16:85,829,569 - 85,830,323 (hg38)

caccg**TGGTGCCCAAGCGTGCCCGG**, aaac**CCGGGCACGCTTGGGCACCA**c (mCherry)

cacc**GAGTCCAGCCTTCAAATCTG**, aaac**CAGATTTGAAGGCTGGACTC** (TagBFP)

*FOS* -3.3kb, chromosome 14:75,274,258 - 75,275,500 (hg38)

caccg**TGTATAAAGAACACCCCAG,** aaac**CTGGGGTGTTCTTTATACA**c (mCherry)

caccg**TAAAAAGTGGAGCTCACACA**, aaac**TGTGTGAGCTCCACTTTTTA**c (TagBFP)

*FOS* +8.9kb, chromosome 14:75,287,748 - 75,287,973 (hg38)

caccg**ATAGGGTACATTGAATCCTG**,aaac**CAGGATTCAATGTACCCTAT**c (mCherry)

cacc**GAGCTGATGGCCATAAGGCC**, aaac**GGCCTTATGGCCATCAGCTC** (TagBFP)

For transfection, plasmids were prepared using an Invitrogen Plasmid Midiprep Kit (Invitrogen, K210004). Transfection of BLAER cells was carried out by nucleofection (Amaxa Nucleofector, Lonza) using Kit C and program X-001, following the manufacturer’s protocol. For each cell line, simultaneous transfection of px330_mCherry and px330_TagBFP was performed in the above combinations. We used 3 million cells per nucleofection reaction alongside 3 micrograms of each plasmid. Cells were cultured for two days after nucleofection. Alive cells that were mCherry+ and TagBFP+ were then sorted on a FACSAria II flow cytometer (BD Biosciences). Single cells were seeded in 96-well plates and were left to grow for two weeks.

Following two weeks of incubation, visible colonies were transferred to a new 96-well plate. A mirror 96-well plate was then established in order to extract genomic DNA from each well and determine which wells harbor cells with the desired excisions. We centrifuged the mirror plates, removed excess medium, and added 50 ul of QuickExtract DNA Extraction Solution (Biosearch Technologies, QE09050) per well. Cell lysates were transferred to PCR 96-well plates and samples were heated in a thermal cycler at 65°C for 10 minutes, followed by heating at 98°C for 5 minutes. Final lysates were diluted by 100 ul of ddH20 per well. 4ul of each well was used for a PCR reaction.

### PCR screening

We performed PCR reactions using a Phusion High-Fidelity DNA Polymerase (ThermoFisher, F530L) to screen for potential homozygous enhancer deletions. PCR primers for each reaction were designed to be at least 100bp upstream and downstream of predicted CAS9 cuts. The following primer combinations were used:

*IRF8* +83kb

CCTGGAGCAGTGATGGACTC (forward)

AGACCTCCTTGCAGAACAGC (reverse)

*IRF8* -69kb

GAAGTGGTTCCATCCGCCT (forward)

AACATCACTCCAGAGAGCCCA (reverse)

*IRF8* DKO (Derivative of *IRF8* -69kb KO clone, +83kb guides used):

CCTGGAGCAGTGATGGACTC (forward)

AGACCTCCTTGCAGAACAGC (reverse)

*IRF8* DKO (Derivative of *IRF8* +83kb KO clone, -69kb guides used):

GAAGTGGTTCCATCCGCCT (forward)

AACATCACTCCAGAGAGCCCA (reverse)

*FOS* -3.3kb

TGTGAGTCCCACAGGAATTG (forward)

CAATCGAGCTTACAGGGTAGC (reverse)

*FOS* +8.9kb

ATTGTCTGTCTCTTATCCCTGAACT (forward)

TTGGACCACTCTGCTAAATTGGAT (reverse)

*FOS* DKO (Derivative of *Fos* +8.9kb KO clone, -3.3kb guides used):

TGTGAGTCCCACAGGAATTG (forward)

CAATCGAGCTTACAGGGTAGC (reverse)

### Flow cytometry

To monitor transdifferentiation, we stained surface markers CD19 and MAC1 and performed flow cytometry. Briefly, cells were collected, washed with PBS, and spun down. Cells were resuspended in a blocking solution for 10 min at room temperature using a human FcR binding inhibitor (1:10 dilution, eBiosciences, catalog no. 16–9161–73). Cells were subsequently stained with antibodies against CD19 (APC-Cy7 mouse anti-human CD19, BD Pharmingen, catalog no. 557791) and MAC1 (APC mouse anti-human CD11b/Mac1, BD Pharmingen, catalog no. 550019) at 4 °C for 20 minutes in the dark. Cells were afterward washed with PBS and resuspended in PBS with DAPI. FACS analysis was carried out in an LSRII instrument (BD Biosciences). Collected data was analyzed using FlowJo software.

### RNA extraction, reverse transcription and qPCR

RNA was extracted with RNeasy Mini Kit (Qiagen, 74104) following the manufacturer’s protocol. Extracted RNA was quantified using a NanoDrop spectrophotometer. 500ng of extracted RNA was used as input for a reverse transcription reaction using the High-Capacity cDNA Reverse Transcription Kit (Applied Biosystems, 4368814). cDNA was then used for quantitative PCR with SYBR Green QPCR Master Mix (Applied Biosystems, A25742). Three technical replicates per gene and condition were used. The following primers, spanning exon junctions, were used to determine gene expression levels of *IRF8, FOS* and *GUSB* (housekeeping gene):

*IRF8*

AGGAGCCTTCTGTGGACGAT (forward)

ACCATCTGGGAGAATGCTGA (reverse)

*FOS*

CTGTCAACGCGCAGGACTT (forward)

GCAGTGACCGTGGGAATGAA (reverse)

*GUSB*

CACCAGGATCCACCTCTGAT (forward)

TCCAAATGAGCTCTCCAACC (reverse)

### Gene expression modeling

Expression values were normalized to the average expression of WT cells within the same transdifferentiation batch. Normalized expression values were used to fit generalized linear models described by Dukler et al., 2017. A brief explanation of the models can be found below:

Additive model: Enhancers control the expression of a gene in an independent, linear fashion. Gene expression can be modeled as:

Expression = b_0_ + b_1_*x_1_ + b_2_*x_2_ + ε

With x_1_ and x_2_ being binary variables describing whether an enhancer is present or excised, b_0_ an intercept term, b_1_, and b_2_ coefficients describing enhancer activity and ε an error term capturing biological and technical noise.

Exponential/Synergistic model: Enhancers control expression multiplicatively, as expected if enhancers display synergistic behavior. Gene expression is modeled as:

Expression = e ^b0 + b1*x1 + b2*x2^ + ε

Logistic model: Transcription of a gene is associated with a low-energy state, while a baseline high-energy state inhibits transcription. Enhancer activity modulates the fraction of time the gene is in its low-energy state:

Expression = g / (1+ e ^−(b0 + b1*x1 + b2*x2)^) + ε

The term g represents the maximum expression level of a gene.

Model coefficients were fit in R using the superEnhancerModelR package (Dukler et al., 2017).

## Acknowledgments

We would like to thank the Graf and Nikolaou lab members for helpful discussions about the project, as well as Sergi Aranda and Sergi Cuartero for reading the manuscript before submission. We also thank members of the Flow Cytometry and Advanced Light Microscopy core facilities. This study was supported by CRG internal funds and the Spanish Ministry of Economy, Industry and Competitiveness (MEIC) Plan Estatal 2019 with project reference number PID2019-109354GB-I00.

