## Supplementary Figures for "Synergistic and antagonistic activities of *IRF8* and *FOS* enhancer pairs during an immune cell fate switch"

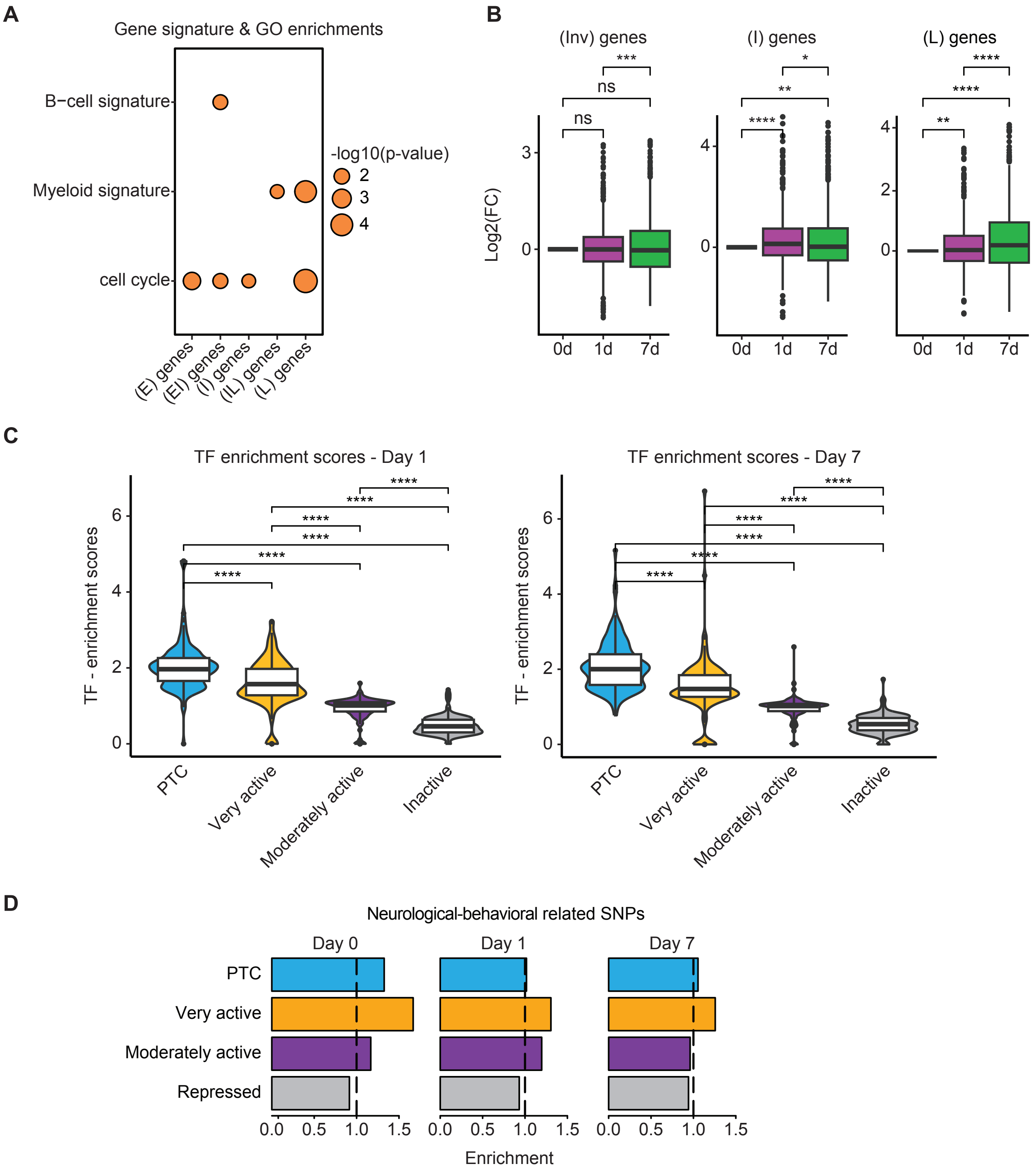
A
Gene signature & GO enrichments
B−cell signature
Myeloid signature
cell cycle
-log10(p-value)
2
3
4
(E) genes
(EI) genes
(I) genes
(IL) genes
(L) genes
B
(Inv) genes
(I) genes
(L) genes
Log2(FC)
ns
***
**
*
****
0d
1d
7d
C
TF enrichment scores - Day 1
TF enrichment scores - Day 7
TF - enrichment scores
****
PTC
Very active
Moderately active
Inactive
D
Neurological-behavioral related SNPs
Day 0
Day 1
Day 7
PTC
Very active
Moderately active
Repressed
0.0
0.5
1.0
1.5
Enrichment

**Supplementary Figure 1. PTCs contain highly expressed, lineage-specific genes across all transdifferentiation stages.**

A. Cell signature enrichments of genes within PTC clusters. A custom list of B-cell-specific and myeloid-specific genes was downloaded from (Monaco et al., 2019). Overlap significance between the generated gene lists and the genes within each PTC cluster was assessed via a hypergeometric distribution test. Gene ontology (GO) enrichment analysis for the genes of each PTC group was performed using gprofiler2 (Kolberg et al., 2020) with the "cell cycle" enriched GO term being depicted. P-values are represented as circles, with the circle area being proportional to the p-value.

B. Expression dynamics of genes falling in the "Invariant", "Intermediate" and "Late" PTC clusters. Values were processed as in Fig.1D. The Wilcoxon signed-rank test was used to determine statistically significant differences (ns p-value > 0.05; * p-value <= 0.05; ** p-value <= 0.01; *** p-value <= 0.001; **** p-value <= 0.0001).

C. Transcription factor-target enrichment scores for the four SEGCOND genomic groups at Days 1 and 7 of transdifferentiation. Scores were calculated as in Fig. 1F. Statistically significant differences were determined via a Wilcoxon rank-sum test (ns p-value > 0.05; * p-value <= 0.05; ** p-value <= 0.01; *** p-value <= 0.001; **** p-value <= 0.0001).

D. Overlap enrichment of SEGCOND genomic categories with neurological-associated single nucleotide polymorphisms (SNPs) as in Fig.1H. No significant overlap was observed with this SNP category.

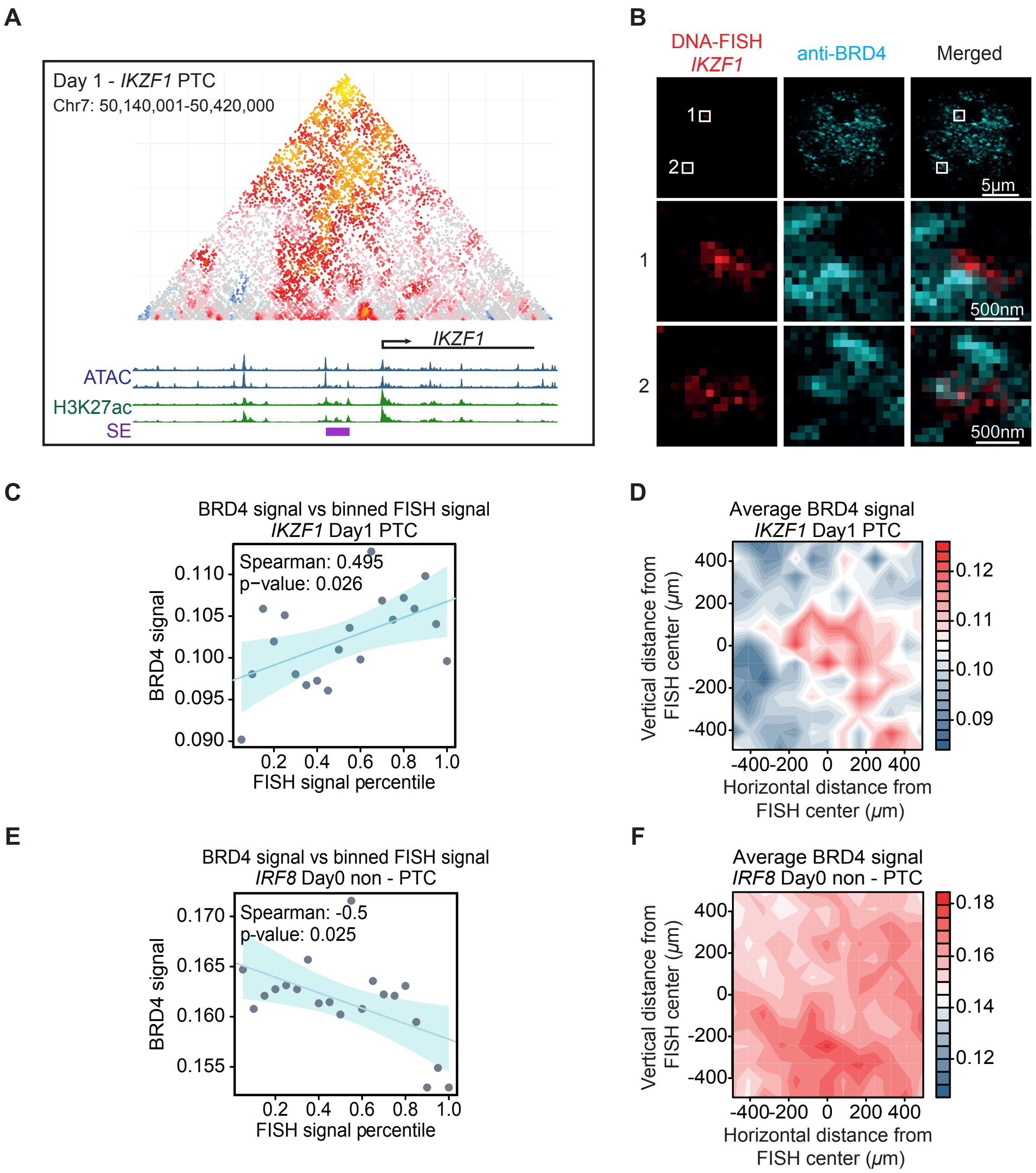

A
Day 1 - IKZF1 PTC
Chr7: 50,140,001-50,420,000
IKZF1
ATAC
H3K27ac
SE
B
DNA-FISH
IKZF1
anti-BRD4
Merged
1
2
5μm
500nm
500nm
C
BRD4 signal vs binned FISH signal
IKZF1 Day1 PTC
Spearman: 0.495
p−value: 0.026
BRD4 signal
0.110
0.105
0.100
0.095
0.090
0.2 0.4 0.6 0.8 1.0
FISH signal percentile
D
Average BRD4 signal
IKZF1 Day1 PTC
Vertical distance from FISH center (μm)
400
200
0
-200
-400
-400-200 0 200 400
Horizontal distance from FISH center (μm)
0.12
0.11
0.10
0.09
E
BRD4 signal vs binned FISH signal
IRF8 Day0 non - PTC
Spearman: -0.5
p-value: 0.025
BRD4 signal
0.170
0.165
0.160
0.155
0.2 0.4 0.6 0.8 1.0
FISH signal percentile
F
Average BRD4 signal
IRF8 Day0 non - PTC
Vertical distance from FISH center (μm)
400
200
0
-200
-400
-400-200 0 200 400
Horizontal distance from FISH center (μm)
0.18
0.16
0.14
0.12

**Supplementary Figure 2. BRD4 hubs associate with active PTCs.**

A. Overview of the *IKZF1* PTC at Day 1 cells. As in Fig.2A.

B. DNA-FISH coupled with BRD4 immunofluorescence, targeting the *IKZF1* PTC at Day 1 cells. The third and second rows of images correspond to the zoomed regions falling within the first and second highlighted areas (white frames).

C. DNA-FISH and BRD4 signal correlation analysis for Day1 *IKZF1* PTC data. Same as in Fig. 2C

D. Contour plots of BRD4 signal enrichment over *IKZF1* Day 1 DNA-FISH centers. Same as in Fig. 2D.

E. DNA-FISH and BRD4 signal correlation analysis for Day 0 *IRF8* PTC data. Same as in Fig. 2C. Higher DNA-FISH values anti-correlate with high BRD4 values, indicating that BRD4 doesn't associate with the *IRF8* PTC at Day 0 cells.

F. Contour plots of BRD4 signal enrichment over *IRF8* Day 0 DNA-FISH centers. Same as in Fig. 2D. No enrichment of BRD4 signal at DNA-FISH centers, marking the *IRF8* PTC, can be observed at Day 0 cells.

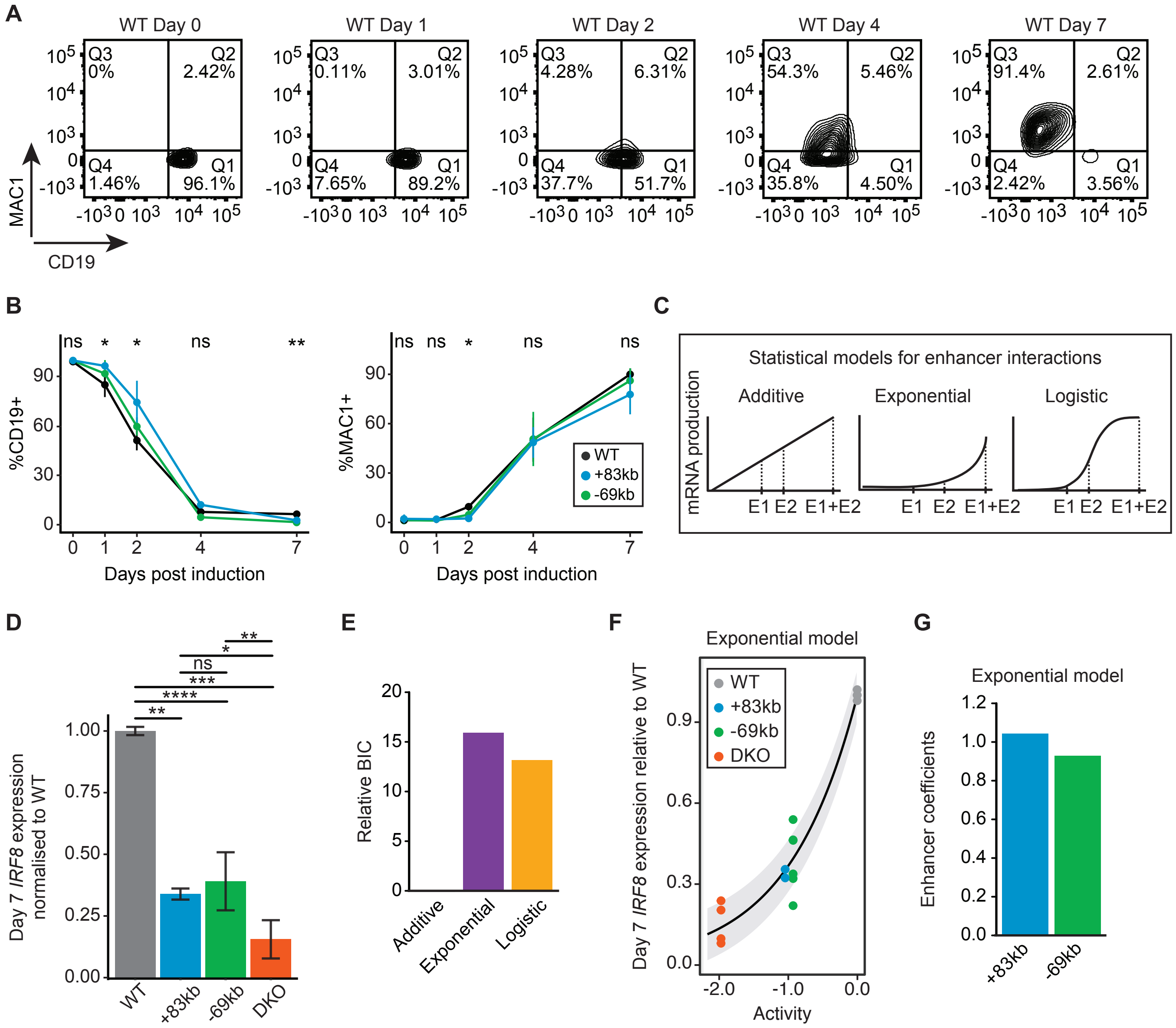

A
WT Day 0
WT Day 1
WT Day 2
WT Day 4
WT Day 7
Q3 0%
Q2 2.42%
Q4 1.46%
Q1 96.1%
Q3 0.11%
Q2 3.01%
Q4 7.65%
Q1 89.2%
Q3 4.28%
Q2 6.31%
Q4 37.7%
Q1 51.7%
Q3 54.3%
Q2 5.46%
Q4 35.8%
Q1 4.50%
Q3 91.4%
Q2 2.61%
Q4 2.42%
Q1 3.56%
MAC1
CD19
B
%CD19+
%MAC1+
Days post induction
ns * * ns **
ns ns * ns ns
WT
+83kb
-69kb
C
Statistical models for enhancer interactions
Additive
Exponential
Logistic
mRNA production
E1 E2 E1+E2
D
Day 7 IRF8 expression normalised to WT
WT
+83kb
-69kb
DKO
E
Relative BIC
Additive
Exponential
Logistic
F
Exponential model
Day 7 IRF8 expression relative to WT
Activity
WT
+83kb
-69kb
DKO
G
Exponential model
Enhancer coefficients
+83kb
-69kb

**Supplementary Figure 3. The -69kb and +83kb *IRF8* enhancers synergize to drive high *IRF8* expression levels.**

A. FACS gating strategy used to determine CD19+ and MAC1+ cells during transdifferentiation. Cells falling within Q1 & Q2 were deemed as CD19+, whereas cells within Q3 & Q4 as MAC1+.

B. Transdifferentiation kinetics of *IRF8* -69kb KO and +83KO cells compared to BLAER cells, monitored via FACS. The mean ± s.d is depicted for each timepoint. Statistical significance was determined using a one-way ANOVA test (ns p-value > 0.05; * p-value <= 0.05; ** p-value <= 0.01; *** p-value <= 0.001; **** p-value <= 0.0001).

C. Overview of statistical models adapted from (Dukler et al., 2016). In brief, the additive model assumes that each enhancer independently adds its activity, linearly increasing gene expression levels. The exponential model assumes synergy between enhancers, leading to exponential increases in gene expression levels. Finally. the logistic model predicts that transcription occurs in a low-energy state, with each enhancer independently reducing the energy threshold required to reach it.

D. Day 7 *IRF8* expression levels of all generated KO lines and WT cells. Same as Fig. 3D.

E. Evaluation of best statistical model fit for Day 7 *IRF8* expression data. As in Fig. 3E, except for the exponential model being the best fit for the dataset.

F. Exponential model fit to *IRF8* Day 7 expression data. Same as in Fig. 3F.

G. Exponential model enhancer coefficients for *IRF8* Day 7 expression data. As in Fig. 3G, except that the model assigns very similar activities to the two enhancers.

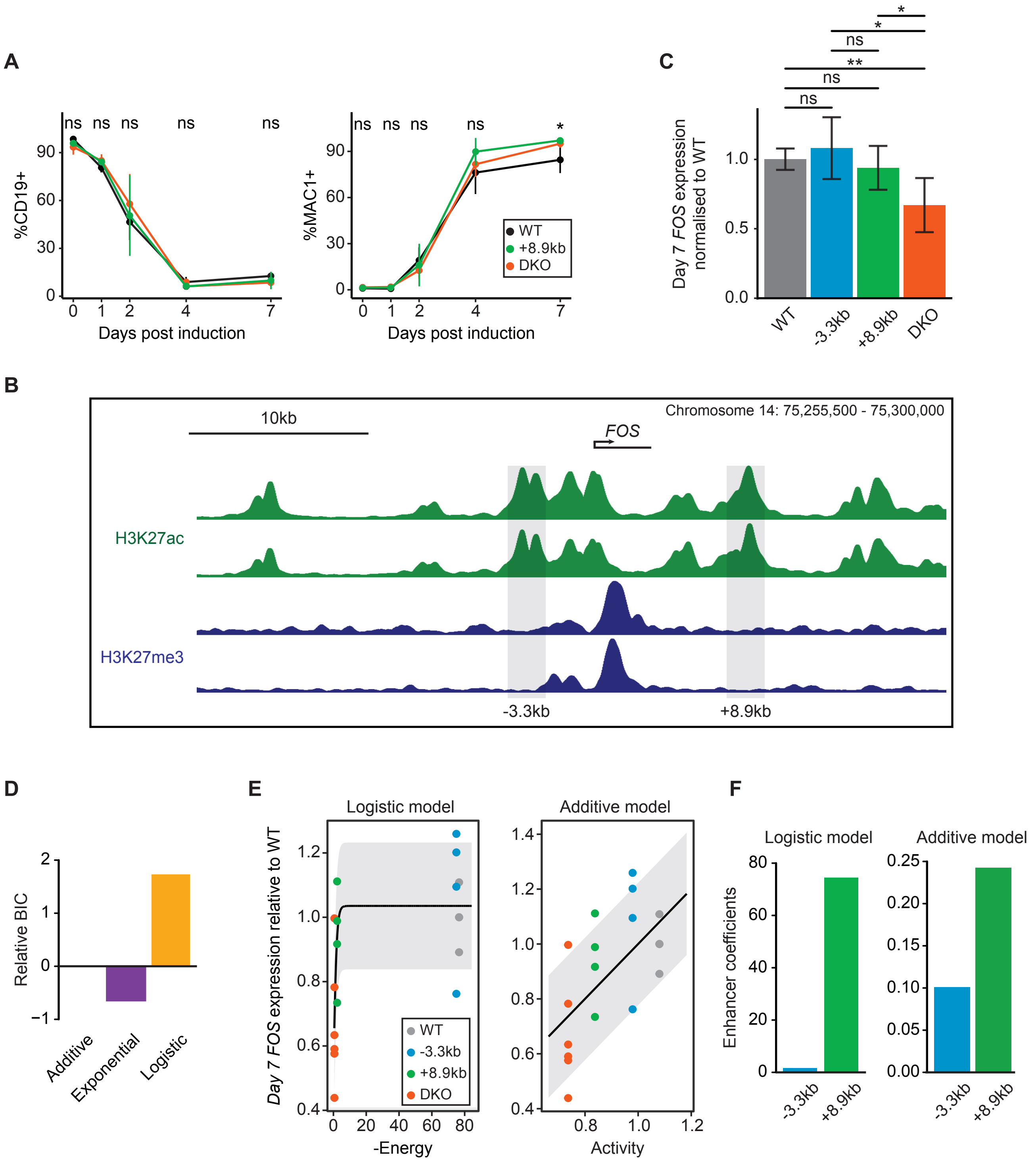
A
%CD19+
Days post induction
%MAC1+
Days post induction
WT
+8.9kb
DKO
ns
*
B
10kb
Chromosome 14: 75,255,500 - 75,300,000
FOS
H3K27ac
H3K27me3
-3.3kb
+8.9kb
C
Day 7 FOS expression normalised to WT
WT
-3.3kb
+8.9kb
DKO
D
Relative BIC
Additive
Exponential
Logistic
E
Logistic model
Additive model
Day 7 FOS expression relative to WT
-Energy
Activity
F
Logistic model
Additive model
Enhancer coefficients

**Supplementary Figure 4. The -3.3kb and +8.9kb *FOS* enhancers cooperate at Day 7 cells, safeguarding *FOS* expression.**

A. Transdifferentiation kinetics of *FOS* +8.9kb KO and DKO cells compared to BLAER cells, monitored via FACS. The mean ± s.d of CD19+ and MAC1+ cells is shown. Statistical significance was determined using a paired Student's t-test (ns p-value > 0.05; * p-value <= 0.05; ** p-value <= 0.01; *** p-value <= 0.001; **** p-value <= 0.0001).

B. Overview of the H3K27ac and H3K27me3 marks in the *FOS* PTC at Day 1 cells. Two H3K27ac and two H3K27me3 ChIP-seq tracks of two biological replicates are depicted. The -3.3kb enhancer isn't decorated with the repressive histone mark H3K27me3.

C. Day 7 *FOS* expression levels of KO and WT cells. As in Fig. 4C. Statistical significance was determined using a Student's t-test (ns p-value > 0.05; * p-value <= 0.05; ** p-value <= 0.01; *** p-value <= 0.001; **** p-value <= 0.0001).

D. Evaluation of best model fit for Day 7 *FOS* expression data. As in Fig. 3E. Both the logistic and additive models were deemed valid.

E. Logistic and additive model fits to *FOS* Day 7 expression data. Same as in Fig. 3F.

F. Logistic and additive model enhancer coefficients for Day 7 *FOS* expression data. As in Fig. 3F. The +8.9kb enhancer activity is proposed to be higher than the activity of the -3.3kb enhancer.
